# Water quality assessments and metagenomic analysis of the polluted river Apatlaco, Mexico

**DOI:** 10.1101/536128

**Authors:** Luz Breton-Deval, Katy Juárez, Alejandro Sánchez-Flores, Rosario Vera-Estrella

**Author notes:** **Corresponding author:** Dr. Luz Breton-Deval, Instituto de Biotecnología, Universidad, Nacional Autónoma de México, Avenida, Universidad 2001, Colonia Chamilpa, Cuernavaca, Morelos 62210, México.

## Abstract

The aim of this study is to analyze the water quality parameters and bacterial diversity and thereby understand the effect of water quality on the microbial population structure in the river. The following parameters: total coliforms, chemical oxygen demand, harness, ammonium, nitrite, nitrate, total Kjeldahl nitrogen, dissolved oxygen, total phosphorus, total dissolved solids, and temperature were analyzing along 17 sampling points in the river. The worst levels of pollution were 510 mg/L chemical oxygen demand, 7 mg/L nitrite, 45 mg/L nitrate, 2 mg/L dissolved oxygen, and 756 mg/L of total dissolved solids. Whole metagenome shotgun sequencing was performed at 4 key points along the river (P1,P7,P10 and P17), the first point had clean water and the other points were polluted, as a result of this pollution, the structure of microbial communities along the river have changed. *Proteobacteria* and *Bacteroidetes* were the most representative phyla with a relative abundance of 57 and 43% respectively for P1, 82 and 15% for P7, 69 and 27% for P10 and 87 and 10% for the last point P17. P1 is rich in microorganism such as *Limnohabitans* a planktonic bacterium very common in freshwater ecosystems. However, in P7, P10 and P17 are rich in opportunistic pathogens such as *Acinetobacter Arcobacter and Myroides* that endangers the health of around 1.6 million people which live around the area. These results elucidate the influence of the pollution on the microbial community and the likely effects on the health of the people around.

## 1. Introduction

Microbial communities in river systems are diverse and dynamic in composition, sometimes as a consequence of environmental changes, diverse stresses and varying nutrient profiles [1]. Microbial communities participate in carbon nitrogen and other biogeochemical cycles in water as well as driving primary production through photosynthetic activity [2]. Furthermore, the maintenance and biodegradation activities are linked to microbial diversity of superficial waters [3]. In this case study, the discharge of wastewater from industrial and urban activities, perhaps has been impacted the structure and function of microbial communities and has consequently disrupted the nutrient cycles and negatively affected energy flow to higher trophic levels [4,5]. Furthermore, contaminants can represent a dangerous source of pathogens for the people who live within the river basin [6]. Rodríguez-Tapia et al [7] developed an ecological model which correlates epidemiological studies with the pollution present in The Atoyac River located in Puebla, Mexico, finding a causal relationship between water contamination and gastrointestinal diseases. This highlights how important it is for communities to have access to clean water for hydration, food production and sanitation. If proper sanitation services are not available, rates of disease are further increased by polluted water [8].

The metabolic potential of superficial water microorganisms is crucial for the transformation and degradation of the pollutants present, including heavy metals, pesticides, and organic waste. Therefore, understanding the diversity and structure of microbial communities can help to indicate the level of pollution in the system and allow us to anticipate the rehabilitation of aquatic ecosystems [9].

The current study was conducted in The Apatlaco River, a waterway located in Morelos, Mexico. This river receives 321 official wastewater discharges, 158 are from the industrial sector and 137 and 26 are from the domestic and agricultural sectors, respectively. Therefore, the pollutants present in The Apatlaco River are very diverse in type and high in volume (CONAGUA-SEMARNAT, 2016). At the origin of the river (P1) the water is clean, and the river doesn’t receive any discharge. Later parts of the river receive numerous different kinds of discharges and consequently the water quality is similar to wastewater. As a result, P1 provides a clean water source to which we have been able to compare measurements from other parts of the river.

Scientific knowledge about the diversity, function, and metabolic pathways of microorganisms in polluted environments is necessary. The Microorganisms present in waterways and the identification of chemical reactions linked to metabolism present in polluted environments can significantly contribute to providing scientific evidence of the effect of pollutants on microbiome diversity and their role as a selective pressure.

In the past, studies of ecology and diversity of microbes were approached with classical techniques. However, most of the microorganism in the environment are unculturable [10]. Currently, advances in DNA sequencing technology such as whole metagenome shotgun (WMS) exhibits a rich profile of microbial communities due to all DNA is cut into tiny fragments and sequenced, resolution is broader than the universal 16S marker gene profiles, which only analyze the 16S locus [11]. The aim of this study is to analyze water quality parameters and bacterial diversity and thereby understand the effect of water quality on the microbial population structure in the river.

## 2. Materials and Methods

### 2.1 Study site and sample collection

The Apatlaco River passes through the small state of Morelos in central Mexico (Figure 1). The Apatlaco River is 63 km long and runs through 10 municipalities in Morelos. Due to its short length and its water’s heterogeneous characteristics, the river provides a convenient model for the evaluation of anthropogenic effects on microbiota. In Morelos there are two primary seasons: the rainy season and the dry season. The rainy season lasts from June to September and during the rest of the year the climate is approximately 24 ± 2 °C. The dry season is the best time to conduct field research given that the microorganisms present during the rainy season are not representative of stable river conditions. Water samples were collected at 17 sampling points along The Apatlaco River (table 1). Sampling field trips were conducted every two months during 2018 in order to span the entire cycle of hydrological variation. Methanogenic analyses used samples from four key points of the river: P1, P7, P10 and P17. At each point (P1, P7, P10, P17), 10 different water samples were collected and then combined to form a single sample for molecular analyses.

**Table 1.**
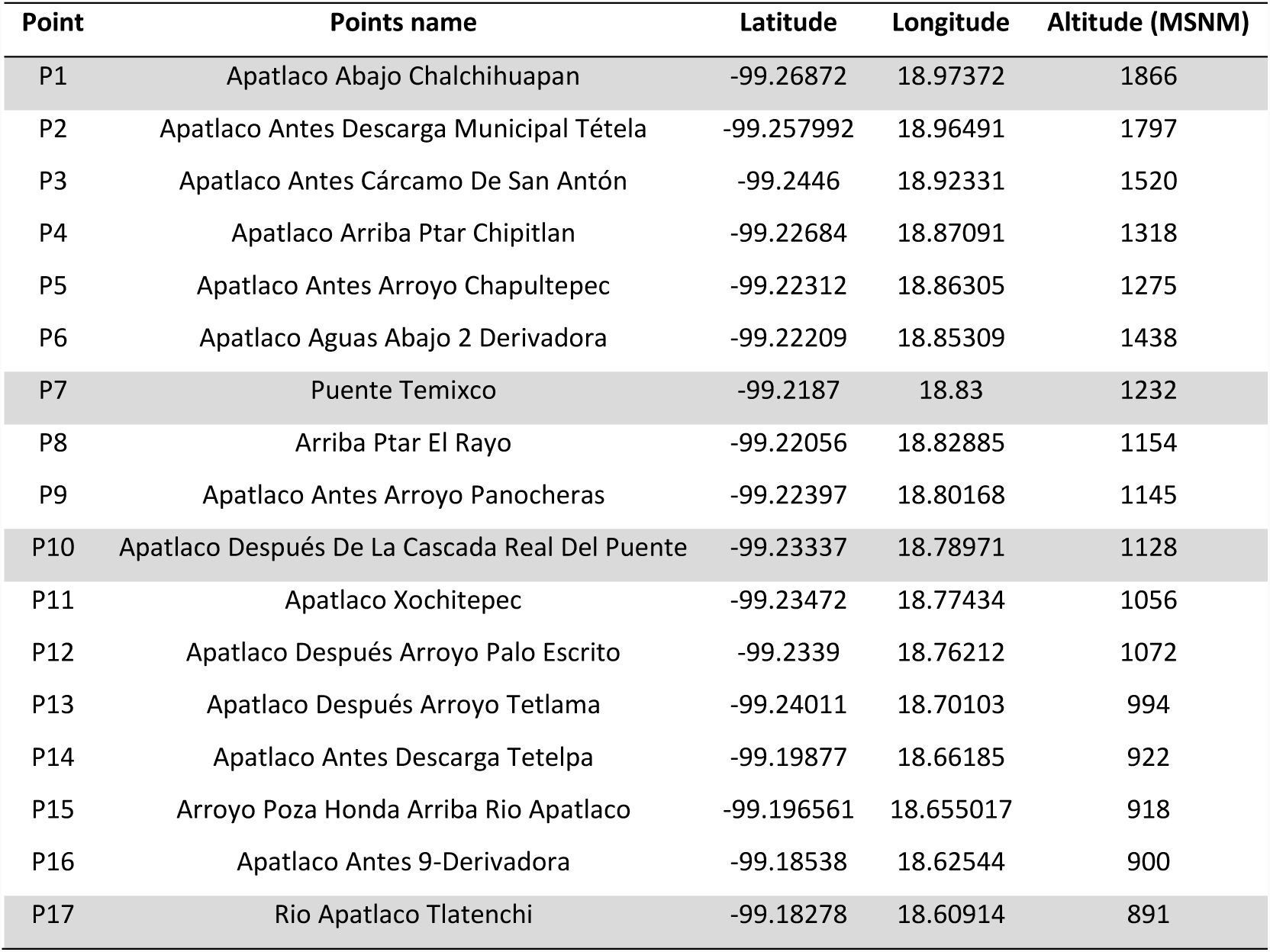
Sampling points in the Apatlaco River

**Figure 1.**
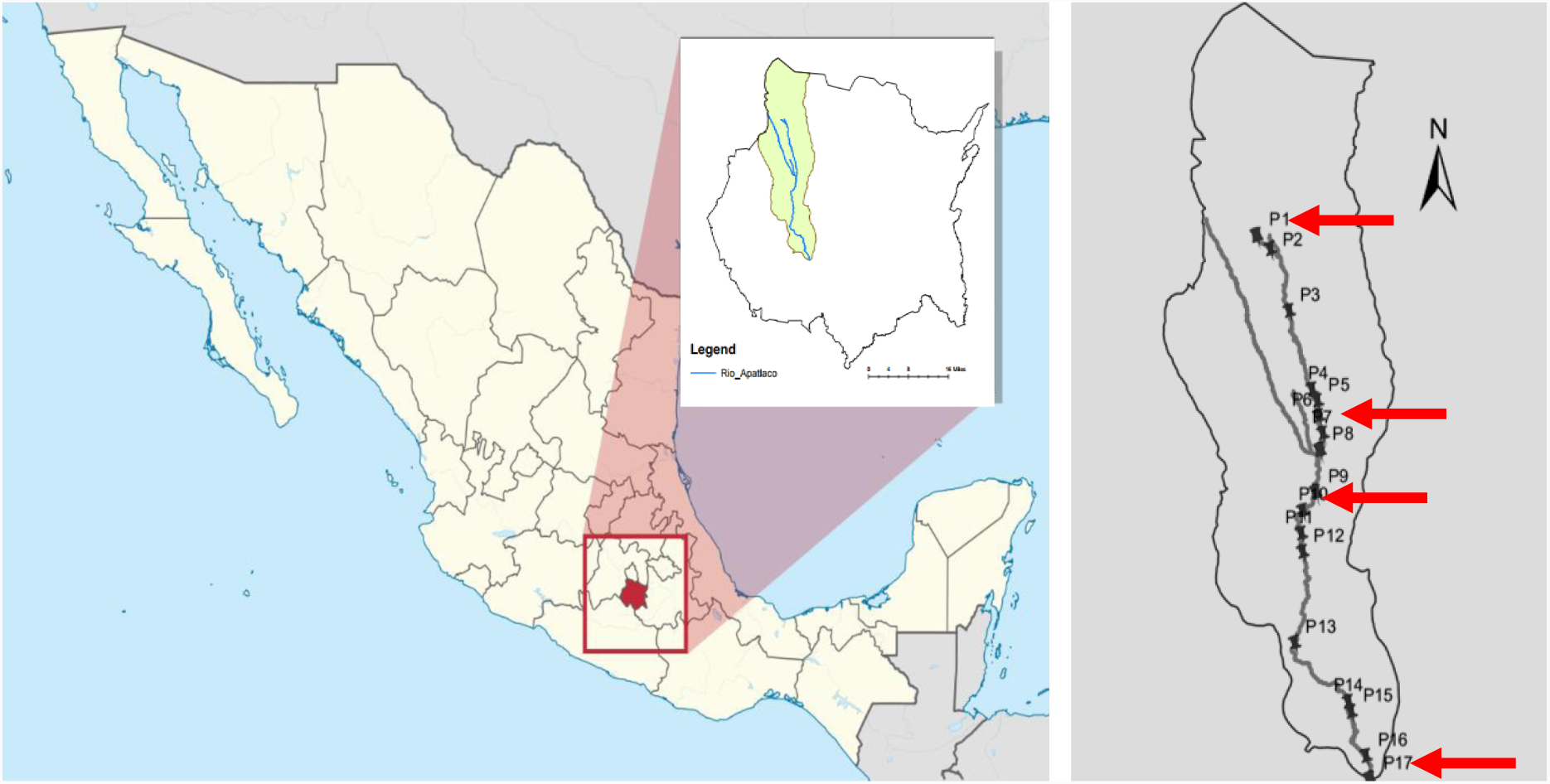
The Apatlaco River basin with the names of sampling points along the river. The red arrows indicate the sampling points (P1, P7, P10 and P17) used for the metagenomic analysis

### 2.2 Environmental parameters measured in the Apatlaco River

Physical and chemical parameters of the river were measured on site using a HANNA multi parametric instrument DR900. This device measures key parameters such as pH, dissolved oxygen (OD), turbidity, ammonium (NH), nitrite (NO2-N) nitrate (NO3-N). Total nitrogen (TN), total phosphorus (TP) and chemical oxygen demand (COD) were determined with the colorimetric method using HACH digester DRB200. Temperature was measured by a thermometer, Total coliforms and total dissolved solids (TDS) were measured according to Standard Methods [12]

### 2.3 DNA extraction and Sequencing

DNA was extracted from water samples using a DNeasy PowerWater Kit (QIAGEN, Hilden, Germany). For each sample, an Illumina library was prepared from total DNA using the TruSeq kit v2 (Illumina, Inc., San Diego, CA, USA) following the manufacturer’s specifications with an average fragment size of 500 bp. The sequencing was performed on the NextSeq500 (Illumina, Inc., San Diego, CA, USA) platform with a 150-cycle configuration, generating paired-end reads with a length of 75 bp.

### 2.4 Data Analysis

After performing a quality control analysis using the FASTQC program [13], the taxonomic profiling was performed using the raw reads with the software MetaPhlan v2.0 [14] using the following parameters: --bt2_ps sensitive-local --min_alignment_len 95 --input_type fastq.

In addition to the MetaPhlan analysis, the software MEGAN6 was used to evaluate the taxonomic distribution of each read based on the comparison of BLASTX with all the sequences in the NCBI-NR databank.

### 2.5 Statistical analysis

Statgraphics Centurion Software (Statpoint Technologies, Inc., Warrenton, Virginia, USA) was used for the principal component analysis (PCA) and to measure the water quality differences between the polluted and unpolluted sites (ANOVA).

## 3. Results and Discussion

### 3.1 Water quality in The Apatlaco River

At the first sample point (P1) the physicochemical analysis showed values of water quality suitable for irrigation, fishing, and recreation. The water quality of the river at this location is good and the aquatic ecosystem is healthy (Table 2, Figure 2) [15,16]. However, wastewater effluents start to enter the river and contaminate the water from the second sample point (P2) onwards. This contamination can be observed via the increased presence of fecal coliforms, collections of microorganisms that live in the intestines of humans and animals, at higher rates than the legally permitted limits. At P2, the river has levels of 200 coliform colonies/100 ml and therefore this water cannot be used for recreational activities, nor fishing or irrigation. The most significant presence is at P7, where 31400 coliform colonies/100 ml are present. Another important parameter widely used to examine organic pollutants in water systems is chemical oxygen demand (COD). The highest values of COD were located at P5 with 320 mg/L and P12 with 360 mg/L as a result of 137 domestic discharge points. Typical wastewater has a COD around 400 mg/l [17]. The rate of COD also affects the values of DO; high values of COD cause low levels of DO. DO values follow the same trend found for COD; the worst levels of DO were found between P5 and P12 at around 3 mg/L. Low levels of DO can cause anoxic zones in the river and this stimulates microorganisms such as sulfate-reducing bacteria that release noxious gases such as hydrogen sulfide, or methanogenic bacteria that release methane gases; one of the most potent greenhouse gases [18].

**Table 2.**
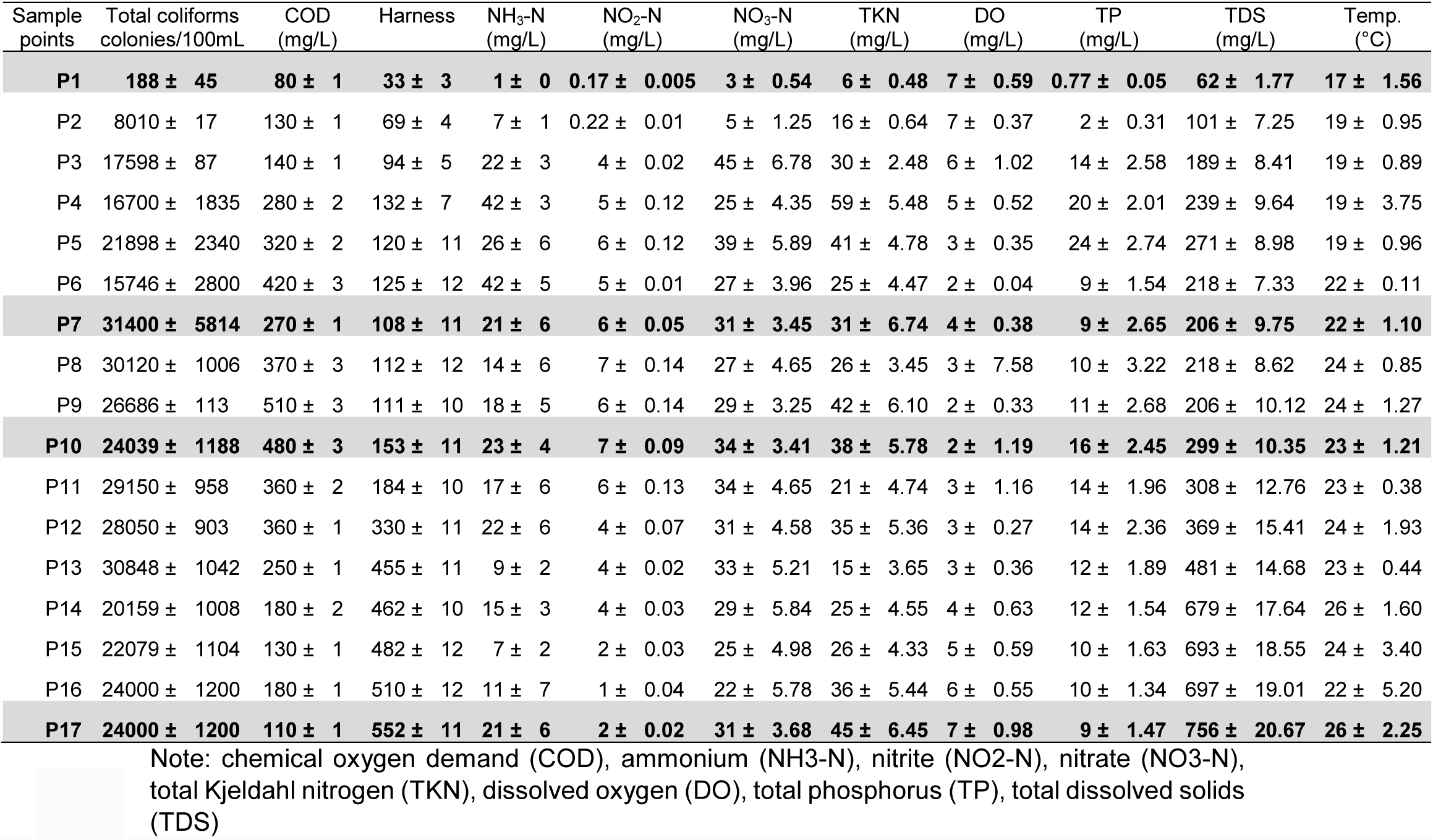
The measurements of chemical parameters along the river

| Sample points | Total coliforms colonies/100mL | COD (mg/L) | Harness | NH<sub>3</sub>-N (mg/L) | NO<sub>2</sub>-N (mg/L) | NO<sub>3</sub>-N (mg/L) | TKN (mg/L) | DO (mg/L) | TP (mg/L) | TDS (mg/L) | Temp. (°C) |
| --- | --- | --- | --- | --- | --- | --- | --- | --- | --- | --- | --- |
| **P1** | **188 ± 45** | **80 ± 1** | **33 ± 3** | **1 ± 0** | **0.17 ± 0.005** | **3 ± 0.54** | **6 ± 0.48** | **7 ± 0.59** | **0.77 ± 0.05** | **62 ± 1.77** | **17 ± 1.56** |
| P2 | 8010 ± 17 | 130 ± 1 | 69 ± 4 | 7 ± 1 | 0.22 ± 0.01 | 5 ± 1.25 | 16 ± 0.64 | 7 ± 0.37 | 2 ± 0.31 | 101 ± 7.25 | 19 ± 0.95 |
| P3 | 17598 ± 87 | 140 ± 1 | 94 ± 5 | 22 ± 3 | 4 ± 0.02 | 45 ± 6.78 | 30 ± 2.48 | 6 ± 1.02 | 14 ± 2.58 | 189 ± 8.41 | 19 ± 0.89 |
| P4 | 16700 ± 1835 | 280 ± 2 | 132 ± 7 | 42 ± 3 | 5 ± 0.12 | 25 ± 4.35 | 59 ± 5.48 | 5 ± 0.52 | 20 ± 2.01 | 239 ± 9.64 | 19 ± 3.75 |
| P5 | 21898 ± 2340 | 320 ± 2 | 120 ± 11 | 26 ± 6 | 6 ± 0.12 | 39 ± 5.89 | 41 ± 4.78 | 3 ± 0.35 | 24 ± 2.74 | 271 ± 8.98 | 19 ± 0.96 |
| P6 | 15746 ± 2800 | 420 ± 3 | 125 ± 12 | 42 ± 5 | 5 ± 0.01 | 27 ± 3.96 | 25 ± 4.47 | 2 ± 0.04 | 9 ± 1.54 | 218 ± 7.33 | 22 ± 0.11 |
| **P7** | **31400 ± 5814** | **270 ± 1** | **108 ± 11** | **21 ± 6** | **6 ± 0.05** | **31 ± 3.45** | **31 ± 6.74** | **4 ± 0.38** | **9 ± 2.65** | **206 ± 9.75** | **22 ± 1.10** |
| P8 | 30120 ± 1006 | 370 ± 3 | 112 ± 12 | 14 ± 6 | 7 ± 0.14 | 27 ± 4.65 | 26 ± 3.45 | 3 ± 7.58 | 10 ± 3.22 | 218 ± 8.62 | 24 ± 0.85 |
| P9 | 26686 ± 113 | 510 ± 3 | 111 ± 10 | 18 ± 5 | 6 ± 0.14 | 29 ± 3.25 | 42 ± 6.10 | 2 ± 0.33 | 11 ± 2.68 | 206 ± 10.12 | 24 ± 1.27 |
| **P10** | **24039 ± 1188** | **480 ± 3** | **153 ± 11** | **23 ± 4** | **7 ± 0.09** | **34 ± 3.41** | **38 ± 5.78** | **2 ± 1.19** | **16 ± 2.45** | **299 ± 10.35** | **23 ± 1.21** |
| P11 | 29150 ± 958 | 360 ± 2 | 184 ± 10 | 17 ± 6 | 6 ± 0.13 | 34 ± 4.65 | 21 ± 4.74 | 3 ± 1.16 | 14 ± 1.96 | 308 ± 12.76 | 23 ± 0.38 |
| P12 | 28050 ± 903 | 360 ± 1 | 330 ± 11 | 22 ± 6 | 4 ± 0.07 | 31 ± 4.58 | 35 ± 5.36 | 3 ± 0.27 | 14 ± 2.36 | 369 ± 15.41 | 24 ± 1.93 |
| P13 | 30848 ± 1042 | 250 ± 1 | 455 ± 11 | 9 ± 2 | 4 ± 0.02 | 33 ± 5.21 | 15 ± 3.65 | 3 ± 0.36 | 12 ± 1.89 | 481 ± 14.68 | 23 ± 0.44 |
| P14 | 20159 ± 1008 | 180 ± 2 | 462 ± 10 | 15 ± 3 | 4 ± 0.03 | 29 ± 5.84 | 25 ± 4.55 | 4 ± 0.63 | 12 ± 1.54 | 679 ± 17.64 | 26 ± 1.60 |
| P15 | 22079 ± 1104 | 130 ± 1 | 482 ± 12 | 7 ± 2 | 2 ± 0.03 | 25 ± 4.98 | 26 ± 4.33 | 5 ± 0.59 | 10 ± 1.63 | 693 ± 18.55 | 24 ± 3.40 |
| P16 | 24000 ± 1200 | 180 ± 1 | 510 ± 12 | 11 ± 7 | 1 ± 0.04 | 22 ± 5.78 | 36 ± 5.44 | 6 ± 0.55 | 10 ± 1.34 | 697 ± 19.01 | 22 ± 5.20 |
| **P17** | **24000 ± 1200** | **110 ± 1** | **552 ± 11** | **21 ± 6** | **2 ± 0.02** | **31 ± 3.68** | **45 ± 6.45** | **7 ± 0.98** | **9 ± 1.47** | **756 ± 20.67** | **26 ± 2.25** |

Note: chemical oxygen demand (COD), ammonium (NH<sub>3</sub>-N), nitrite (NO<sub>2</sub>-N), nitrate (NO<sub>3</sub>-N), total Kjeldahl nitrogen (TKN), dissolved oxygen (DO), total phosphorus (TP), total dissolved solids (TDS)

**Figure 2.**
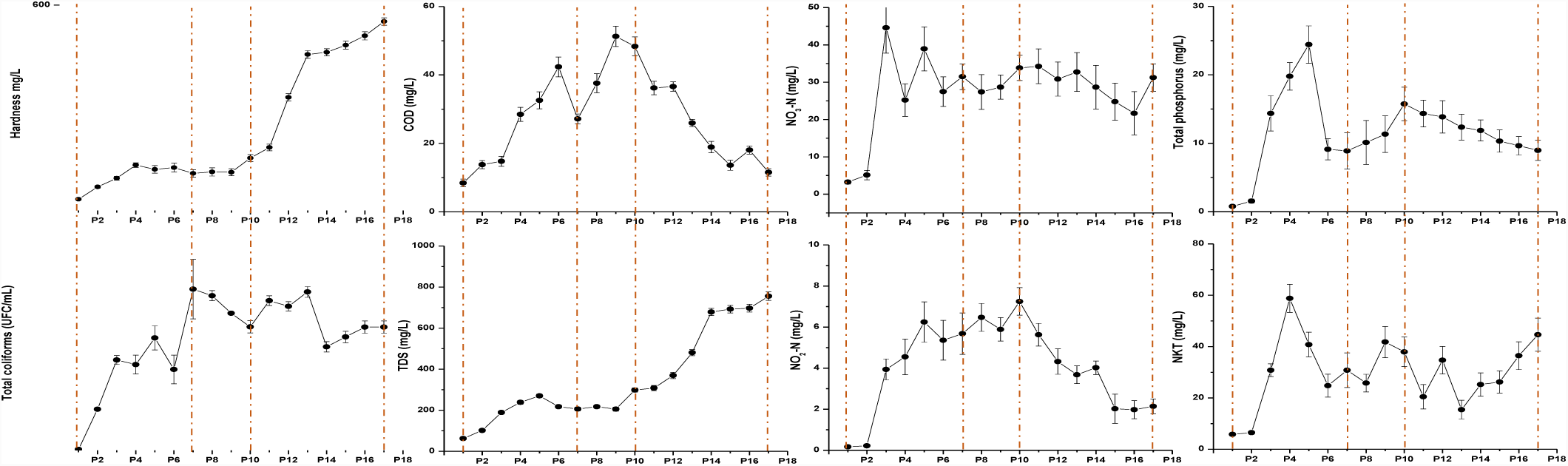
The most representative chemical parameters along the river. The red dotted lines indicate the sampling points (P1, P7, P10 and P17) used for the metagenomic analysis

The total dissolved solids (TDS) and the degree of hardness both increase over the course of the river due to the accumulation of ions and other compounds such as calcium, chloride, sodium, nitrates, phosphates and metals derived from the dumping of industrial wastewater [19].

The concentration of different species of nitrogen is correlated with season and river flow. However, there are some rivers in which one of these factors, flow or season, is more influential than the other. For example, the total nitrogen and nitrate collected in The Stillwater River and The Great Miami River (both in Fairfield, Ohio, USA) tends to increase as the current intensifies. In contrast, the same parameters in The Mad River in Dayton, Ohio, tend to remain constant for the entire length of the river and throughout the year [20]. Regarding the different species of nitrogen in The Apatlaco River, the NO_2_-N values start to increase from P3 onwards, with values of 4 mg/L. It is well known that levels up to 1 mg/L of NO_2_-N can react with proteins and form carcinogenic N-nitrosamines [21]. The same trend can be observed for NO_3_-N, where abnormal values appear at P3, with 45 mg/L. The source of NO_3_-N could be organic or inorganic and possibly come from waste discharge, animal waste or artificial fertilizers. Both anions (NO_2_ and NO_3_) were found in lesser quantities from P13 onwards, however the factor that most affects the concentration of nitrogen species along this river is not the different flow rates but the multiple discharges of wastewater along the length of this body water. This effect is more evident with the values of NH_3_-N and TKN that showed fluctuations possibly associated with discharges of wastewater [22]. The concentration of total phosphorus starts at 0.77 mg/L and varies along the river, the highest point (24 mg/L) is reached at sampling point 5.

### 3.2 Taxonomic assignment of metagenomic data along four sites in The Apatlaco River

Taking in account variations of physicochemical parameters along the waterway were selected four sites along the river in order to perform the metagenomic analysis. Figure 3 provides a map of bacteria diversity in the different sampling points as can be seen the diversity in every point is different. The principal phyla in terms of relative abundance at P1 were *Proteobacteria* and *Bacteroidetes* with 57% and 43%, respectively. 51% of the *Proteobacteria* phyla corresponded to *Betaproteobacteria* class and *Burkholderiales* accounted 50% of total at order level, 47% of which were *Comamonadacea*, 44% were classified as *Limnohabitans* genus*. Limnohabitans* is a common member of bacterioplankton in freshwater habitats which plays an important role in carbon transfer to higher trophic levels [23–25]. Another important freshwater planktonic bacterium is *Polynucleobacter* which is usually found in the same habitats as *Limnohabitans* [26]. At P7 the level of *Limnohabitans* decreased to 15% which is strong evidence of the severe impact of wastewater contamination of the freshwater ecosystem. At P10, where there is a discharge of the local PTAR, the relative abundance of *Limnohabitans* declines further to 5%. It is possible that the bacteria from the PTAR discharge compete with *Limnohabitans* and thereby displace them. P7 is severely impacted by wastewater discharges from both industrial and domestic source. At this site, *Proteobacteria* was the most representative Phulym with 82% abundance and *Bacteroidetes* decreased to 15 %. At Class level the abundance was: *Gammaproteobateria* 53%, *Betaproteobacteria* 25%, *Epsilonproteobacteria* 3%, *Flavobacteria* 12% and *Sphingobacteria* 3%. The *Pseudomonadales* 52% order was the most abundant order within *Gammaproteobacteria*, which included the *Moraxellaceae* family 50% and *Pseudomonadaceae* family 2%. *Acinetobacter* is represented with 49% and it is important to mention that because this genera are free living saprophytes widely used in biotechnological applications including bioremediation of several compounds such as phenol, hexadecane, diesel etc. but also some species as *A. baumanii* is a water organism recently identified as an opportunistic pathogen in humans, affecting people with compromised immune systems, and is becoming increasingly important as a hospital-derived (nosocomial) infection [27,28]. The other family identified was *Pseudomonadaceae.* The principal genera of *Pseudomonadaceae* was *Pseudomonas* 1.7%, a microorganism with the ability to adapt and colonize several ecological environments including water, sewage, soil, plants, animals, and humans [29]. Their powerful metabolism facilitates their frequent use in bioremediation projects as it is well known that they can degrade phenols, hydrocarbons, pesticides, heavy metals and dyes [30–32]. At P1 the relative abundance of *Pseudomonas* was less than 1 (0.7%) however from P7 onwards their relative abundance begins to increase, and at P17 their relative abundance was 7%. The other 15% of the bacteria found at P7 were phylum *Bacteroidetes* where 5% are from the *Sphingobacteriaceae* family and 10% from the *Flavobacteriaceae* family. Compared with P1, the relative abundance of the all genera were diminished. P10 is located following a PTAR treated-water discharge and has the highest levels along the river for parameters such as COD, NH_3_-N and NO_2_-N. (Table 2). In this context, microbial relative abundance changes to 69% for *Proteobacteria* and 27% for *Bacteroidetes.* The most abundant class of *Proteobacteria* phylum was *Epsilonprotobacteria* with 45% of relative abundance, all of which comes from genus *Arcobacter.* Of the 15 species of this genera, *A butzleri, A. cryaerophilus* and *A. skirrowii* are considered emerging pathogens in gastrointestinal infections and extra intestinal invasive diseases [33]. The remaining 25% of relative abundance is dispersed between *Gammaproteobacteria* 15% and *Betaproteobacteria* 9%. Most of the *Gammaproteobacteria* were *Pseudomonadales* 11%, *Moraxellaceae* family with the genera *Acinetobacter* 6% and *Pseudomonadaceae* family with the genera *Pseudomonas* 5%. Almost all *Betaproteobacteria* were *Burkholderiales* 8%.

**Figure 3.**
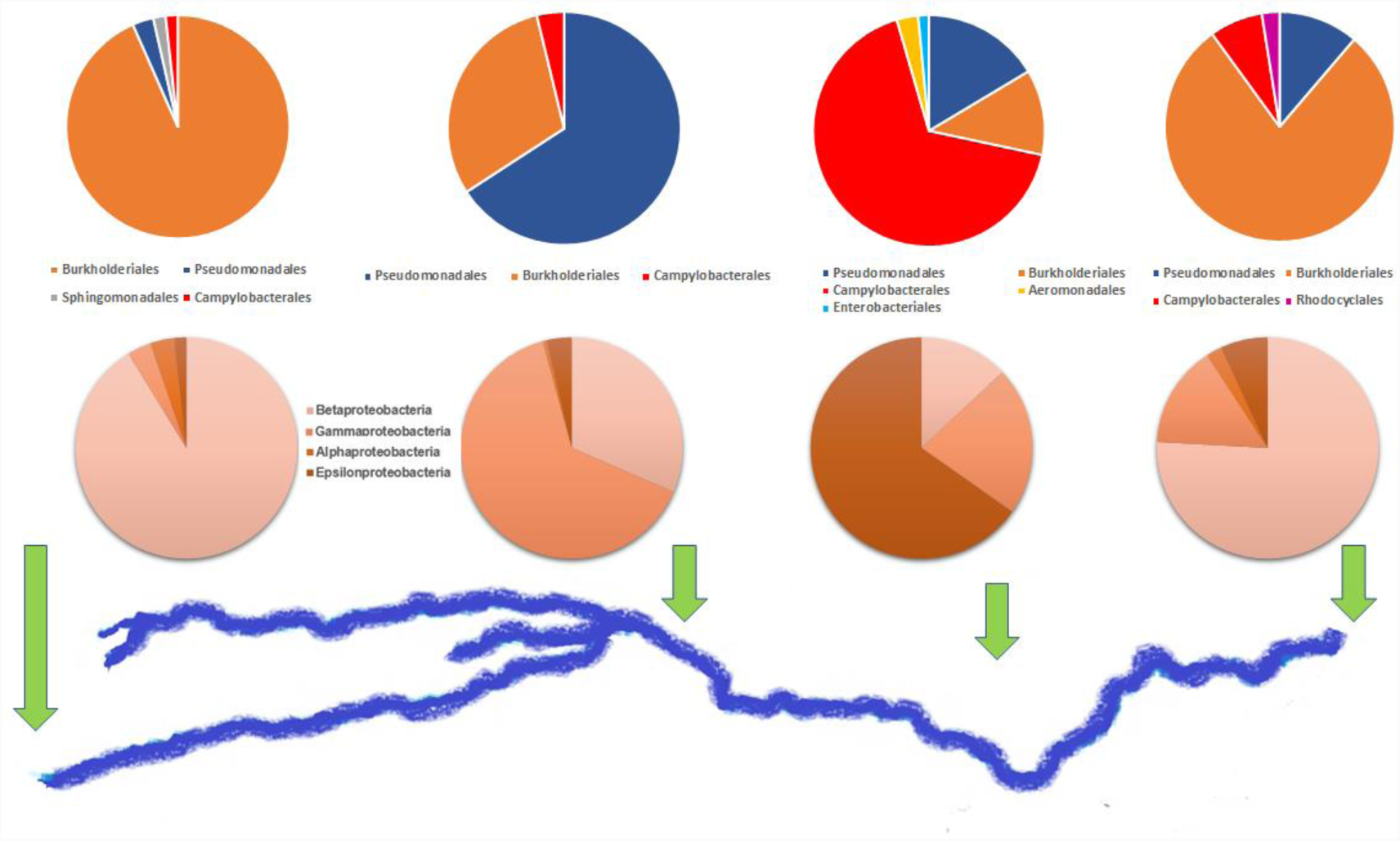
Associations between kind of proteobacteria (down circles) and genera (top circles) with the geographical location in the 4 most relevant points (P1, P7, P10, P17)

While the 27% of the phylum *Bacteroidetes* were distributed in class *Flavobacteriia 15%*, family *Flavobacteriaceae*, mainly in the following genera, *Myroides* 6%, *Riemerella* 4%, *Cellulophaga* 2%, *Gillisia* 1% and 12% in the class *Bacteroidia*, order *Bacteroidales*, families *Prevotellaceae* 7%, *Bacteroidaceae* 4% and *Porphyromonadaceae* 1%.

The final sampling point, P17, present the same trend as P7, an enrichment of *Proteobacteria* 87% and a decrease in *Bacteroidetes* 10%. Most of the *Proteobacteria* were *Betaproteobacteria* class 66% of which 62% were from *Burkholderiales* order, 44% from the *Comamonadaceae* family and 12% from the *Burkholderiaceae* family. The other class were *Gammaproteobacteria* 13%, which *Pseudomonadaceae* genus was 8%, *Flavobacteriia* 7% were Flavobacteriaceae 6%, Epsilonproteobacteria 6% were Campylobacteraceae, *Alphaproteobacteria* 2% and *Actinobacteria* 1% as can be seen in Figure 4.

**Figure 4.**
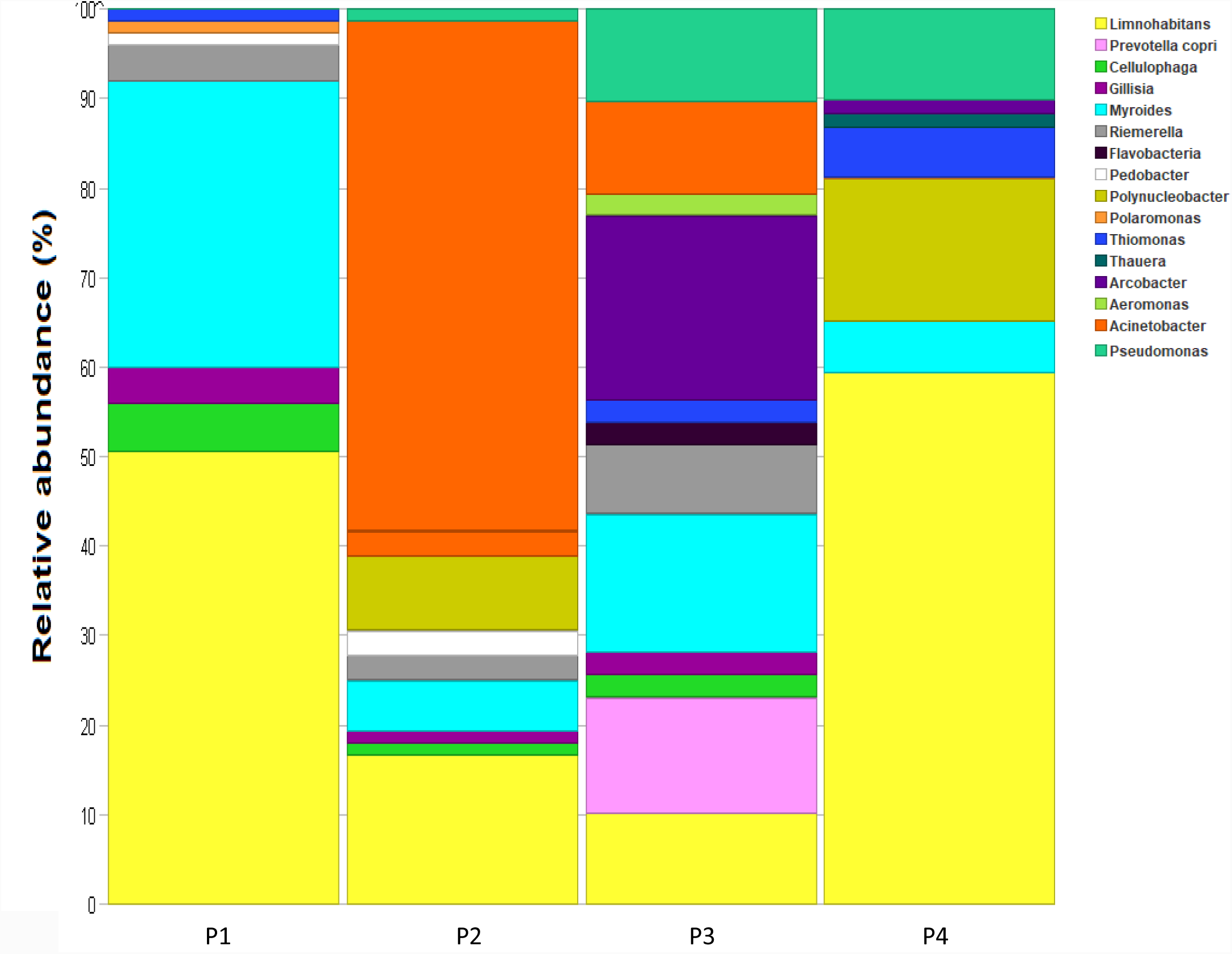
Phylogenetic profile along the river. Where P1 is clean water and P7, P10 and P17 polluted water.

### 3.3 Relationships between water quality factors and bacterial community

The corresponding ANOVA indicates that polluted sites (P7, P10 y P17) have significant differences (*p = 0.0094*) in parameter levels compared with the unpolluted site, P1, with a significance of 95%.

The PCA showed that the first three principal components explain 87% of the data variance (Table 3). The first PCA explains 46% of the variance and is strongly related (> 0.35) with NO_2_, COD and DO (Figure 5). These water quality parameters describe organic contamination as a result of domestic discharges where high levels of organic matter consume dissolved oxygen present in the river and perhaps due to high amounts of COD, in some parts of the river, there are a simultaneous methanogenesis and nitrogen removal via NO_2_ [34]. As a result, in the P10, the site with highest levels in COD and NO_2_ and the lowest levels in DO the dominant microorganisms who have been able to live with low levels of O_2_ were *Arcobacter* genus which requires around 10% of O_2_ for survive and the genera *Prevotella* which is anaerobic. There are other interesting genus present in every point however some arrived in the discharges and are not a microorganism which live there for the chemical conditions of the water such as *Acinetobacter* genus. This genus has their highest abundance in P7 and in P10 their abundance decreases due to the chemical conditions and the competition with other bacteria. The second PCA explains 30% of the variance and were strongly related (> 0.35) with hardness, TDS and temperature. These water quality parameters provide information about the concentration of dissolved or suspended compounds such as carbonate, chlorides, sulfates, phosphates, nitrates, magnesium, calcium, sodium and metals [35] (fig. 2). The increase in hardness and TDS could block light and thus reduce the rate of photosynthesis, causing a reduction in the levels of dissolved oxygen and altering the trophic chains [36]. The highest levels of TDS, hardness and temperature were at P17. The principal genera founded there were *Limnohabitans*, a planktonic bacterium founded in abundance in P1, is likely the microorganism of this genera tolerates better high levels of TDS and hardness than organic pollution. Other genera that increase their abundance levels in that point were *Thiomonas* which has the ability to oxidize arsenite and thiosulfate and used as energy source [37,38]. As well as, *Pseudomonas* which at that point their abundance showed the highest level. The final PCA explains 12% of the variance and is related to TP and NH_3_, the higher levels of both compounds were between P1 and P7 and after the P10 their levels were constants along the points for that reason is not possible differentiate their impact in the microbial structure.

**Table 3.** Principal component analysis and their weight of the chemical parameters analyzed in the river

| parameters | PC1 | (%) | PC2 | (%) | PC3 | (%) |
| --- | --- | --- | --- | --- | --- | --- |
| COD | 0.38613 | 14.90948 | -0.18010 | 3.24346 | -0.32881 | 10.81180 |
| DO | -0.40753 | 16.60831 | 0.07549 | 0.56986 | 0.31694 | 10.04491 |
| Hardness | -0.02517 | 0.06335 | 0.56230 | 31.61768 | 0.10342 | 1.06949 |
| NH<sub>3</sub> | 0.27547 | 7.58837 | -0.18004 | 3.24151 | 0.44629 | 19.91721 |
| NO<sub>3</sub> | 0.37177 | 13.82144 | 0.12579 | 1.58221 | 0.33589 | 11.28194 |
| NO<sub>2</sub> | 0.43337 | 18.78130 | -0.14377 | 2.06710 | -0.09176 | 0.84205 |
| TCOL | 0.35053 | 12.28678 | 0.26986 | 7.28239 | -0.18528 | 3.43268 |
| TDS | 0.00173 | 0.00030 | 0.55760 | 31.09222 | 0.16103 | 2.59307 |
| TEMP | 0.19851 | 3.94062 | 0.43926 | 19.29520 | -0.34455 | 11.87126 |
| TP | 0.34641 | 12.00013 | -0.00911 | 0.00830 | 0.53043 | 28.13560 |
| Eigenvalue | 4.57900 |  | 2.97440 |  | 1.15430 |  |
| Variability (%) | 45.79100 |  | 29.74400 |  | 11.54300 |  |
| Cumulative | 45.79100 | 100.00 | 75.53500 | 100.00 | 87.07900 | 100.00 |

**Figure 5.**
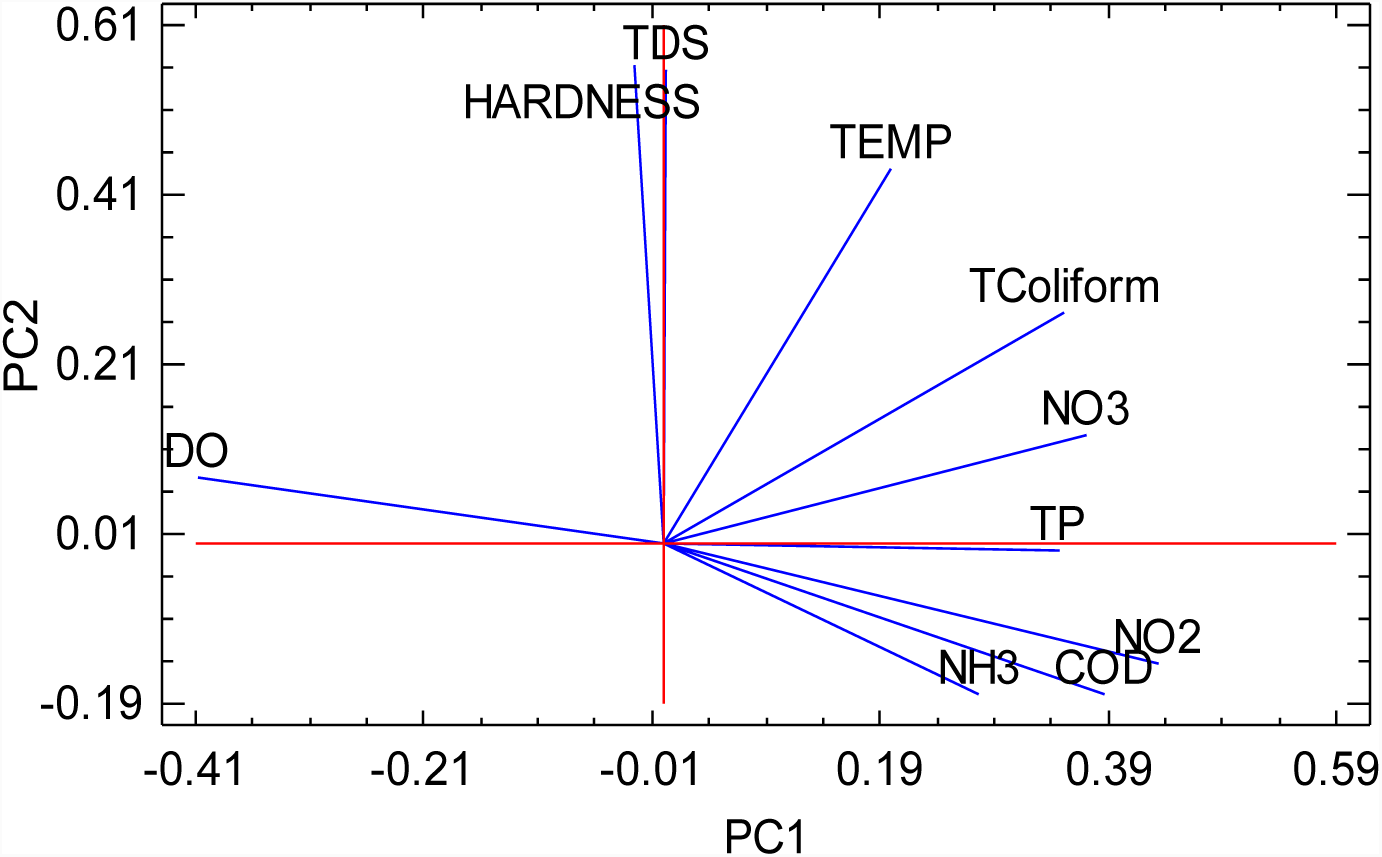
Principal component analysis plot of the chemical parameters

## 4. Conclusion

This study evidence that The River Apatlaco has been damage with different kinds of discharges along their riverbed, P10 has the highest levels in COD, NH_3_, NO_2_, NO_3_, TP and the lowest level in dissolved oxygen while at the end of the river TDS and harness showed their highest levels. The principal component analysis showed that first component explains 46% of the variance in the system and is strongly related with NO_2_, COD and DO, key parameters for develop a healthy aquatic life. As a result of this pollution, the structure of microbial communities along the river have changed. P1 is rich in *Limnohabitans* a planktonic bacterium very common in freshwater ecosystems. However, in P7, P10 and P17 there are microorganism such as *Acinetobacter Arcobacter and Myroides,* which usually are not there and are a danger for human health, now are an important part of the microbial community and this endangers the health of around 1.6 million of people who live in the area around. There are other microorganisms such as *Pseudomonas, Thiomonas* and *Acinetobacter* which have change their abundance due to the changes in the chemical water conditions. These microorganisms have the metabolic machinery to be used in future bioremediation projects if there are stimulate property

## Acknowledgements

The authors thank UNAM-IBT, Mexico for financial support to this research. Also, LB-D thank to Consejo Nacional de Ciencia y Tecnología (CONACYT) and their program CATEDRAS for support the Project 285

